# Latent-Space Residue-Pair Networks from Molecular Dynamics Distinguish Ligand-Activated and SAVI-Associated STING

**DOI:** 10.64898/2026.10.08.757787

**Authors:** Yuko Tsuchiya, Yasushige Yonezawa

**Author notes:** Corresponding author: Yuko Tsuchiya.

## Abstract

Stimulator of interferon genes (STING) provides an informative model system for developing methods to extract and compare residue-pair fluctuation patterns from molecular dynamics (MD) trajectories. We combined a one-dimensional convolutional autoencoder trained on apo STING with temporal mean pooling, Mahalanobis-distance clustering, and residue-pair similarity networks to compare apo, ligand-bound, and SAVI-associated multi-mutant datasets using 25-dimensional latent descriptors. Networks constructed from clusters capturing state-dependent fluctuation patterns showed lid-dominated organization in the ligand-bound dataset and a1b1-enriched, structurally distributed relationships involving residue 281 in the multi-mutant dataset. These findings identify contrasting residue-pair fluctuation signatures in the analyzed trajectories and provide an interpretable framework for generating hypotheses about ligand-associated and mutation-associated dynamic allostery.

## Introduction

Allosteric regulation is a fundamental principle of protein function, whereby perturbations at one site, such as ligand binding or amino-acid substitution, alter the conformational ensemble and functional output at distant sites [1–9]. Despite major advances in molecular dynamics (MD) simulations, extracting mechanistically informative dynamic coupling patterns from high-dimensional trajectories remains challenging, particularly when functionally relevant signals are embedded in a large background of common motions [10–11]. Representation-learning approaches, including autoencoder-based models, can compress high-dimensional MD-derived observables into low-dimensional latent descriptors that retain information about protein dynamics and conformational variation [9,12–15]. Here, we use representation learning to extract compact features from MD-derived residue-pair distance time series, enabling systematic comparison of low-population, state-dependent fluctuation patterns that are difficult to capture with residue-wise summary metrics alone.

A clinically important example of long-range regulation is the innate immune adaptor stimulator of interferon genes (STING) [16]. Upon binding cyclic dinucleotide ligands, STING undergoes conformational rearrangements in its ligand-binding domain (LBD), including closure of a lid region over the ligand-binding pocket [17–18]. These changes propagate toward membrane-proximal connector regions and promote downstream signaling, including C-terminal-tail (CTT) engagement and higher-order assembly [19–20]. In contrast, STING-associated vasculopathy with onset in infancy (SAVI) arises from disease-associated gain-of-function mutations that trigger ligand-independent constitutive signaling [21–22]. Structural and cellular studies indicate that SAVI-associated mutations can perturb autoinhibitory constraints and favor polymerization-competent states [18,20,21–23]. However, how these mutations reorganize residue-pair-level dynamic coupling relative to ligand-induced activation remains incompletely understood. In particular, it is unclear whether mutation-associated constitutive activation simply recapitulates the lid-centered ligand-bound program or instead engages an alternative dynamic coupling regime.

Here, we develop a residue-pair-centric analysis framework and apply it to apo, ligand-bound, and SAVI-associated multi-mutant STING models (Figure 1). Instead of characterizing each residue using a residue-wise summary metric, such as root-mean-square fluctuation, we represent each residue pair by a time series of mean pairwise distances between side-chain heavy atoms along MD trajectories [9,13]. A convolutional autoencoder is trained to compress each 1800-point distance trajectory into a low-dimensional latent representation that captures its fluctuation pattern. Because this representation summarizes temporal behavior rather than static geometry, it can identify residue pairs with similar fluctuation signatures even when the underlying residues are not spatially adjacent.

**Figure 1.**
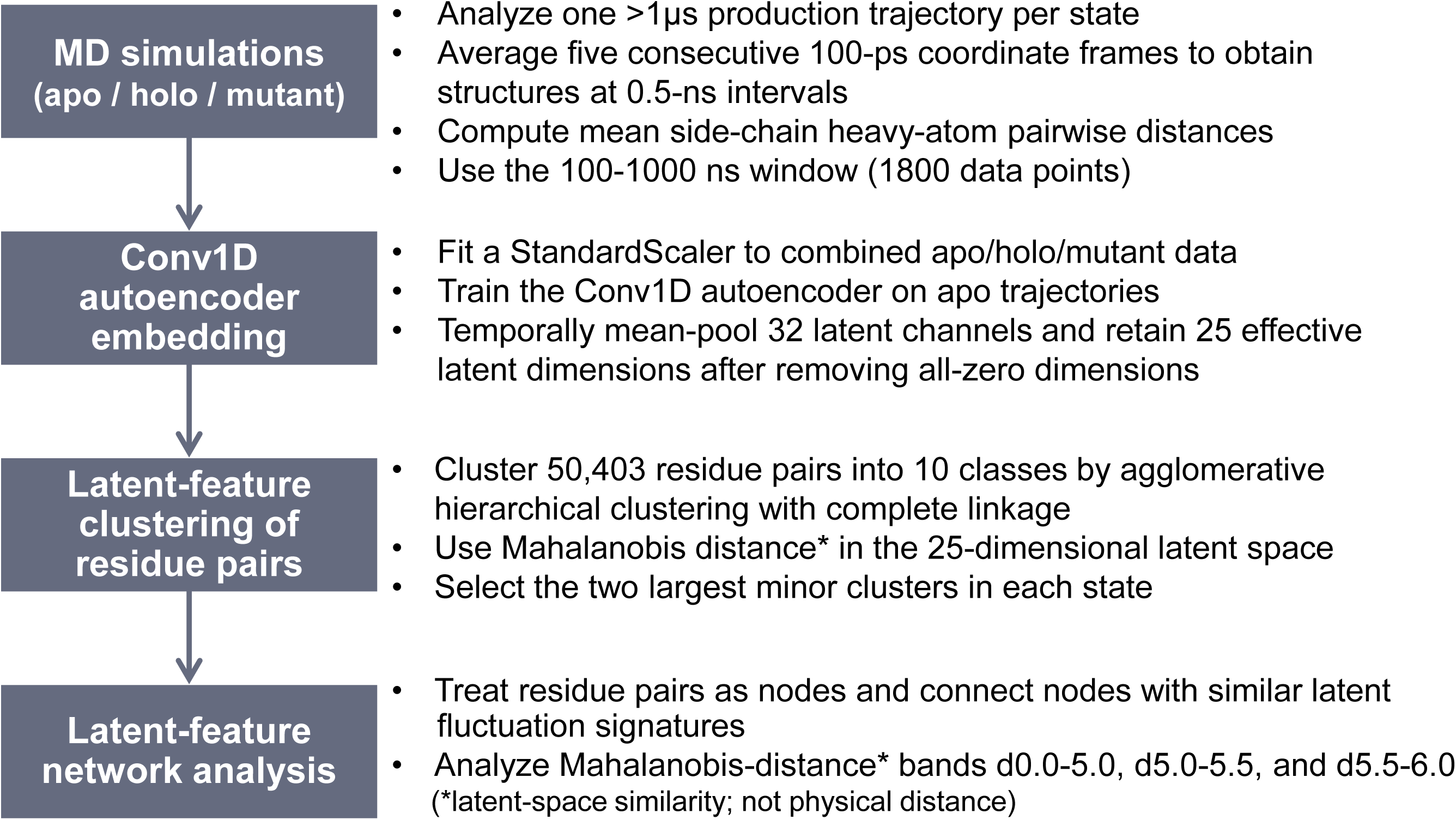
Pipeline overview.

We apply this framework to three STING conditions: apo, ligand-bound (holo), and a SAVI-associated multi-mutant model. Among the mutation sites, we pay particular attention to residues 281 and 284 in the a3 helix of the LBD. Together with residue 206, these positions define a disease-associated region implicated in the control of type I interferon signaling [23], and structural work has linked hyperactive STING mutations to altered CTT sequestration and polymerization competence [20]. Latent vectors are clustered using Mahalanobis distance and agglomerative clustering to isolate low-population residue-pair behaviors. We then construct networks over selected clusters, where nodes correspond to residue pairs and edges denote similarity in latent fluctuation features. By comparing network topology, composition, and centrality across states, we aim to (i) identify which regions dominate coordinated dynamics in each state; (ii) determine how the wild-type counterparts of SAVI hotspot residues, notably 281 and 284, participate in ligand-activated networks; and (iii) characterize mutation-associated network reorganization associated with constitutive-activation-like dynamics.

## Methods

### System preparation and domain definitions

We analyzed STING in three states: apo, holo (ligand-bound), and a SAVI-associated multi-mutant model. The apo and mutant models were based on the human STING structure PDB 6NT5 [18]. The holo model was based on the ligand-bound STING structure PDB 6NT7 [18]. Residue substitutions present in the 6NT7 construct were reverted to the corresponding residues in the apo 6NT5 sequence before simulation.

Residue ranges used to define key regions are summarized in Table 1 and Figure 2A. Each simulation system consisted of a STING dimer (chains PROA and PROB) embedded in a mixed DDPC/DLPE lipid bilayer and solvated with TIP3P water; in the apo setup this corresponded to 308 DDPC, 154 DLPE, 50,362 TIP3 waters, and 148 K+ (POT) / 146 Cl-(CLA) ions, yielding an overall neutral system.

**Figure 2.**
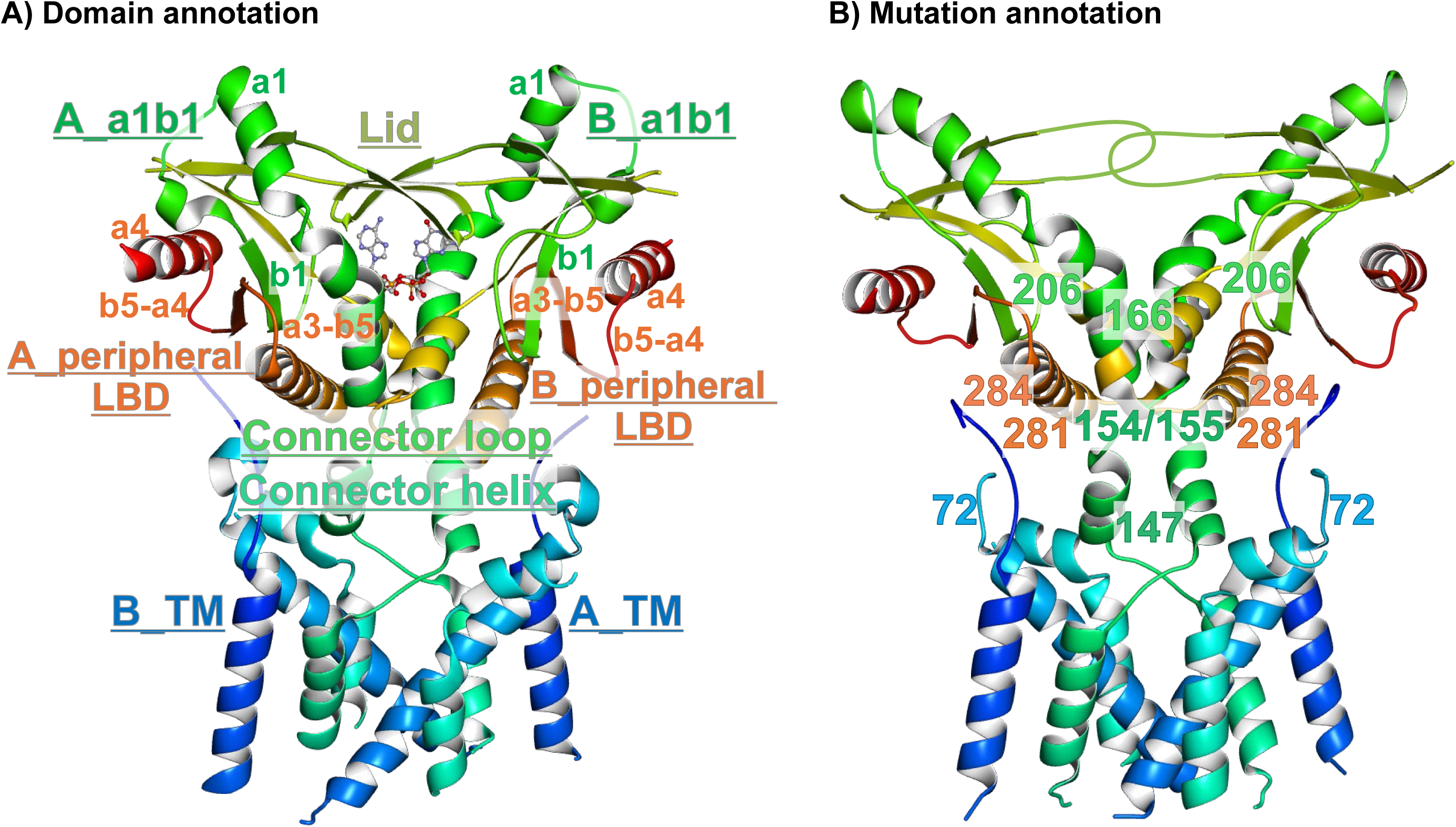
STING dimer structure with domain and mutation annotations. A) Domain annotations: TM (transmembrane), connector helix, connector loop, a1b1, lid, and peripheral-LBD regions depicted on the holo structure. Residue IDs assigned to each region are summarized in Supplementary Table 1. B) Mutation annotations: SAVI-associated mutant residues in the multi-mutant model depicted on the apo structure.

**Table 1.** Domain compositions and clustering summary for the two largest minor clusters used for network analysis.

| Domain composition (#residues) | Cluster ID (#residue pairs) | Apo (130 <sup>1</sup>) |  | Holo (245) |  | Mutant (192) |  |
| --- | --- | --- | --- | --- | --- | --- | --- |
|  |  | Cluster 1 (66 <sup>2</sup>) | Cluster 2 (38 <sup>2</sup>) | Cluster 1 (96) | Cluster 2 (46) | Cluster 1 (82) | Cluster 2 (58) |
| Lid region | 224-244 (13 <sup>3</sup>) | 20 (0.30) | 20 (0.53) | 84 (0.88) | 39 (0.85) | 40 (0.49) | 20 (0.34) |
| Peripheral-LBD region (a3-b5/b5-a4/a4) | 302-308, 315-324, 325-336 (13) | 47 (0.71) | 30 (0.79) | 21 (0.22) | 14 (0.30) | 44 (0.54) | 14 (0.24) |
| a1b1 region | 176-187, 188-197 (10) | 12 (0.18) | 4 (0.11) | 19 (0.20) | 6 (0.13) | 26 (0.32) | 23 (0.40) |
|  | #partner residues in a1-b1 pairs | b5-a4: 9 (0.75)<br>a1-b1: 3 (0.25) | b5-a4: 4 (1.0) | Lid: 15 (0.79)<br>b5-a4: 2 (0.11) | Lid: 5 (0.83)<br>b5-a4: 1 (0.17) | Lid: 9 (0.35)<br>b5-a4: 8 (0.31)<br>a1-b1:6 (0.23) | Lid: 6 (0.26)<br>mutant: 5 (0.22)<br>others: 4 (0.17) |
| #pairs incl. SAVI residues | SAVI residues (8): 72,147,154,155, 166,206,281,284 | SAVI (4):<br>72B_233A<br>72B_234A<br>147A_232A<br>147B_232A | SAVI (2):<br>232A_281A<br>281B_336A | SAVI (6):<br>155A_228A<br>155B_228A<br>189A_284A<br>232A_281A<br>232A_281B<br>233A_284B | SAVI (4):<br>233A_281B<br>233A_284A<br>234A_281B<br>147A_233A | SAVI (1):<br>189B_281B | SAVI (6):<br>191B_281B<br>191B_284B<br>72A_232A<br>147A_233A<br>166A_189A<br>166B_189A |
| #pairs incl. TM residues |  | 22 (0.33) | 3 (0.08) | 8 (0.08) | 6 (0.13) | 19 (0.23) | 26 (0.45) |
| #pairs incl. connector helix/loop |  | 5 (0.08) | 3 (0.08) | 11 (0.11) | 6 (0.13) | 1 (0.01) | 7 (0.12) |

1) Total number of residue pairs outside the numerically dominant background cluster.
2) Numbers of residue pairs in the largest and second-largest minor clusters.
3) Numbers of representative residues assigned to the indicated region.

The mutant model was constructed by introducing eight SAVI-associated substitutions into the apo coordinates: H72N, V147L, N154S, V155M, G166E, C206Y, R281Q, and R284G (Figure 2B). Residues 281 and 284 were highlighted as key SAVI-associated sites because of their reported roles in activation-related interfaces [18,23]. The composite mutant was designed to enrich mutation-associated dynamic signatures and should therefore be interpreted as a state-level stress-test model rather than as a representation of a specific patient genotype. The analysis focuses on the membrane-embedded dimer; explicit CTT engagement and higher-order polymerization are outside the modeled systems.

### Molecular dynamics simulations and pairwise distance trajectories

Force-field setup: Systems were parameterized using a CHARMM-family additive force field converted to GROMACS format by CHARMM-GUI FF Converter (forcefield.itp). Water was modeled with TIP3P (moleculetype TIP3), and ions were represented as K+ (POT) and Cl− (CLA). Nonbonded settings followed CHARMM conventions (combination rule 2, gen-pairs = yes; 1–4 scaling factors fudgeLJ = 1.0 and fudgeQQ = 1.0). Production MD used a 2 fs time step with LINCS constraints on bonds to hydrogen, a Verlet cutoff scheme (neighbor list 1.2 nm), LJ force switching from 1.0 to 1.2 nm, and PME electrostatics with a 1.2 nm real-space cutoff. Temperature was maintained at 303.15 K using a Nose–Hoover thermostat (SOLU, MEMB, SOLV groups; τT = 1 ps), and pressure at 1 bar using a semi-isotropic Parrinello–Rahman barostat (τP = 5 ps; compressibility 4.5×10−5 bar−1).

One production trajectory per state was used for the present analysis. Each trajectory extended beyond 1 μs and was generated with GROMACS on GPU (CUDA); the archived apo production log records gmx mdrun 2022-dev-20220222 (git SHA c07b37733d). Coordinates were written every 100 ps during simulation. Five consecutive saved frames were coordinate-averaged to generate structures at 0.5 ns intervals, and the 100–1000 ns analysis window yielded 1800 structures per state. For each residue pair at each averaged structure, all Euclidean distances between side-chain heavy atoms were calculated and their arithmetic mean was used as the inter-residue distance, producing an 1800-point time series. For glycine, the C *α* atom was used in place of side-chain atoms.

To keep the residue-pair space computationally tractable while retaining inter-regional couplings, we curated the time-series data into 50,403 residue pairs per state. We first selected a representative set of 168 positions, sampled approximately every two to three residues while retaining functionally important sites such as SAVI-associated and lid residues. N- and C-terminal positions outside the analyzed range were then removed (residues ≤17 and ≥337), yielding 159 representative positions. Treating the two protomers separately gave 318 chain-specific residue instances; all unique unordered pairs among these instances were retained, yielding C(318,2) = 50,403 residue pairs (Supplementary Table 1).

To place the three states on a common standardized scale, the residue-pair identifier column was excluded and the apo, holo, and mutant datasets were concatenated along the residue-pair axis. A scikit-learn StandardScaler was fitted to the combined 151,209 × 1,800 matrix, standardizing each of the 1,800 time-point features using the mean and standard deviation across the combined data. The fitted transformation was then applied to each state. Subsequent autoencoder inference, clustering, and network analyses were performed separately for each state using these standardized trajectories.

### Convolutional autoencoder and latent feature extraction

We trained a one-dimensional convolutional autoencoder (Conv1D-AE) on apo-state residue-pair distance trajectories [9,13]. Several architectures were evaluated by varying the number of convolutional layers, filters, and kernel sizes. The selected encoder comprised three Conv1D layers with 256, 128, and 64 filters and kernel sizes of 21, 14, and 7, respectively, followed by a 32-channel latent Conv1D layer (kernel size 1). Encoder convolutions used stride 1, same padding, and ReLU activation, with dropout (0.1) after the first two convolutional layers. The apo dataset was randomly divided into an 80% training portion and a 20% held-out reconstruction subset using random_state = 42; during model fitting, 10% of the training portion was reserved for validation. The batch size was 100, training was performed for 100 epochs, mean-squared error was used as the loss function, and Adam (learning rate 0.001) as the optimizer. Model selection was based on low reconstruction loss on the held-out subset together with visual inspection of representative input and reconstructed distance trajectories; preference was given to models that preserved non-trivial temporal fluctuations and amplitudes rather than producing overly smoothed reconstructions. Because the held-out reconstruction loss contributed to model selection, it was used as a reconstruction criterion rather than as an independent estimate of predictive performance.

For each residue pair, the encoder produced a 32-channel latent feature map over the 1800 time points. Temporal mean pooling across the time axis yielded a 32-dimensional descriptor of the fluctuation pattern. The same trained model was applied to apo, holo, and mutant trajectories. Seven latent dimensions, corresponding to the same columns in all three states, were identically zero and were removed, yielding 25-dimensional effective latent feature vectors for subsequent analyses.

### Clustering of residue-pair fluctuation features

We clustered residue pairs into 10 classes using agglomerative hierarchical clustering with complete linkage on a precomputed Mahalanobis-distance matrix derived from the 25-dimensional latent vectors. For each state, the covariance matrix of the latent vectors was calculated and its pseudoinverse was used in the Mahalanobis-distance calculation. The 10-cluster partition was used to resolve distinct residue-pair fluctuation patterns outside the dominant background. In each state, this procedure yielded one numerically dominant background cluster containing >50,000 pairs and nine smaller clusters with fewer than 100 pairs each, referred to here as minor clusters (Supplementary Table 2). We focused on the two largest minor clusters in each state (Table 1); together, these clusters accounted for more than half of all residue pairs outside the background cluster in every state.

Preliminary comparisons were also performed with K-means and DBSCAN clustering and with Euclidean or cosine-based distances. K-means produced comparatively balanced partitions, whereas DBSCAN assignments were strongly sensitive to density parameters. Agglomerative clustering with Euclidean or cosine distance produced substantially larger non-background partitions. In contrast, agglomerative clustering with Mahalanobis distance separated the numerically dominant background population from multiple low-population clusters, providing the resolution required to compare state-dependent residue-pair fluctuation patterns. We therefore used this combination for the analyses reported below.

### Network construction and quantitative analysis

For network analysis, nodes correspond to residue pairs in the selected minor clusters. Separate networks were constructed by retaining edges whose latent-vector Mahalanobis distances fell within 0.0–5.0, 5.0–5.5, or 5.5–6.0 (denoted d0.0–5.0, d5.0–5.5, and d5.5–6.0, respectively; Table 2 and Supplementary Figure 1). Each network contains the edges in its specified band, rather than all edges below the band’s upper limit. All three networks are derived from the same analyzed time series. The bands distinguish levels of similarity between learned fluctuation descriptors, with smaller distances indicating greater similarity; they do not represent physical inter-residue distances or successive time intervals. Edges thus encode descriptor similarity rather than direct physical contacts.

**Table 2.** Network size and regional composition across clusters and latent-space Mahalanobis-distance bands. A) Apo state. B) Holo state. C) SAVI-associated mutant state. Values report the major connected component used for composition analysis; region categories are non-mutually exclusive.

| Cluster | Distance ranges | #nodes <sup>1</sup> | #edges <sup>1</sup> | regional composition<br>(#pairs including at least one corresponding residue) |  |  | #mutant nodes <sup>4</sup> |
| --- | --- | --- | --- | --- | --- | --- | --- |
|  |  |  |  | a1b1 | lid | peripheral-LBD |  |
| Apo_c1 | d0.0-5.0 | 27<br>(diff <sup>2</sup>:24,<br>same <sup>3</sup>:3) | 91 | 11 (40.7%) | 6 (22.2%) | 21 (77.8%) | [281/284 <sup>5</sup>] 0<br>[others <sup>6</sup>] 2 (147A_232A, 147B_232A)<br>[total <sup>7</sup>] 2 |
|  | d5.0-5.5 | 35<br>(diff:31,<br>same:4) | 84 | 11 (31.4%) | 8 (22.9%) | 29 (82.9%) | [281/284] 0<br>[others] 2 (147A_232A,147B_232A)<br>[total] 2 |
|  | d5.5-6.0 | 42<br>(diff:38,<br>same:4) | 131 | 12 (28.6%) | 9 (21.4%) | 35(83.3%) | [281/284] 0<br>[others] 3 (147A_232A, 147B_232A,<br>72B_233A)<br>[total] 3 |
| Apo_c2 | d0.0-5.0 | 13<br>(diff:11,<br>same:2) | 14 | 2 (15.4%) | 7 (53.8%) | 11 (84.6%) | [281/284] 2 (232A_281A, 281B_336A)<br>[others] 0<br>[total] 2 |
|  | d5.0-5.5 | 17<br>(diff:16,<br>same:1) | 20 | 3 (17.6%) | 8 (47.1%) | 17 (100%) | [281/284] 1 (281B_336A)<br>[others] 0<br>[total] 1 |
|  | d5.5-6.0 | 25<br>(diff:20,<br>same:5) | 51 | 4 (16.0%) | 13 (52.0%) | 23 (92.0%) | [281/284] 2 (232A_281A, 281B_336A)<br>[others] 0<br>[total] 2 |

1) Numbers of nodes and edges in the major connected component used for network-composition analysis.
2) Numbers of nodes composed of residues from different chains.
3) Number of nodes composed of residues from the same chain.
4) Number of nodes containing a residue position associated with a SAVI mutation.
5) Number of nodes containing residue 281 and/or 284 among the mutant nodes.

B) Holo state
| Cluster | Distance ranges | #nodes | #edges | regional composition<br>(#pairs including at least one corresponding residue) |  |  | #mutant nodes |
| --- | --- | --- | --- | --- | --- | --- | --- |
|  |  |  |  | a1b1 | lid | Peripheral-LBD |  |
| Holo_c1 | d0.0-5.0 | 34(diff:26, same:8) | 68 | 5 (14.7%) | 33 (97.1%) | 12 (35.3%) | [281/284] 3 (232A_281A, 232A_281B, 233A_284B)<br>[others] 1 (155B_228A)<br>[total] 4 |
|  | d5.0-5.5 | 38(diff:30, same:8) | 73 | 7 (18.4%) | 37 (97.4%) | 17 (44.7%) | [281/284] 2 (232A_281B, 233A_284B)<br>[others] 1 (155B_228A)<br>[total] 3 |
|  | d5.5-6.0 | 49(diff:36, same:13) | 192 | 11 (22.4%) | 46 (93.9%) | 20 (40.8%) | [281/284] 4 (189A_284A, 232A_281A, 232A_281B, 233A_284B)<br>[others] 1 (155B_228A)<br>[total] 5 |
| Holo_c2 | d0.0-5.0 | 12(diff:9, same:3) | 14 | 0 (0%) | 12 (100%) | 9 (75.0%) | [281/284] 1 (233A_281B)<br>[others] 1 (147A_233A)<br>[total] 2 |
|  | d5.0-5.5 | 15(diff:12, same:3) | 15 | 2 (13.3%) | 14 (93.3%) | 11 (73.3%) | [281/284] 1 (233A_281B)<br>[others] 1 (147A_233A)<br>[total] 2 |
|  | d5.5-6.0 | 20(diff:14, same:6) | 31 | 3 (15.0%) | 19 (95.0%) | 14 (70.0%) | [281/284] 2 (233A_281B, 233A_284A)<br>[others] 1 (147A_233A)<br>[total] 3 |

C) SAVI-associated mutant state
| Cluster | Distance ranges | #nodes | #edges | regional composition<br>(#pairs including at least one<br>corresponding residue) |  |  | #mutant nodes |
| --- | --- | --- | --- | --- | --- | --- | --- |
|  |  |  |  | a1b1 | lid | peripheral-LBD |  |
| Mut_c1 | d0.0-5.0 | 41(diff:38,<br>same:3) | 74 | 19 (46.3%) | 14 (34.1%) | 27 (65.9%) | [total] 0 |
|  | d5.0-5.5 | 45(diff:42,<br>same:3) | 100 | 22 (48.9%) | 16 (35.6%) | 31 (68.9%) | [total] 0 |
|  | d5.5-6.0 | 58(diff:55,<br>same:3) | 195 | 23 (39.7%) | 23 (39.7%) | 40 (69.0%) | [total] 0 |
| Mut_c2 | d0.0-5.0 | 23(diff:14,<br>same:9) | 47 | 14 (60.9%) | 10 (43.5%) | 9 (39.1%) | [281/284] 0<br>[others] 3 (72A_232A, 166A_189A,<br>166B_189A)<br>[total] 3 |
|  | d5.0-5.5 | 29(diff:16,<br>same:13) | 63 | 17 (58.6%) | 12 (41.4%) | 12 (41.4%) | [281/284] 0<br>[others] 3 (72A_232A, 166A_189A,<br>166B_189A)<br>[total] 3 |
|  | d5.5-6.0 | 31(diff:15,<br>same:16) | 101 | 17 (54.8%) | 13 (41.9%) | 12 (38.7%) | [281/284] 1 (191B_281B)<br>[others] 3 (72A_232A, 166A_189A,<br>166B_189A)<br>[total] 4 |

Above a Mahalanobis distance of 6.0, the number of connected node pairs increased steeply and the resulting networks became difficult to interpret as selective similarity relationships. We therefore restricted the network analysis to Mahalanobis distances ≤6.0.

We quantified network size, regional composition, and degree centrality. Regional composition was defined as the fraction of nodes containing at least one residue in each of three non-mutually exclusive regions: a1b1 (the latter half of the a1 helix, residues 176-187, and the loop between the a1 helix and b1 sheet, residues 188-197), lid (residues 224-244, spanning the ligand-binding region between the central portions of the b2 and b4 sheets), and the peripheral LBD region (the a3-b5 loop, residues 302-308; b5-a4 loop, residues 315-324; and a4 helix, residues 325-336) (Table 1). Table 2 reports the major connected component used for network analysis; isolated nodes and small disconnected components were excluded from these counts and figures. Degree centrality was calculated on the full graph before this component filtering, including isolated nodes. For each node, degree centrality was defined as k/(N - 1), where k is its number of incident edges and N is the number of nodes in the full input graph. High-centrality nodes were interpreted as relatively well-connected residue-pair nodes within their respective graphs (Table 3). We also summarized node-type composition and identified subnetworks involving SAVI hotspot residues 281 and/or 284 (Table 2 and Supplementary Figure 1).

**Table 3.**
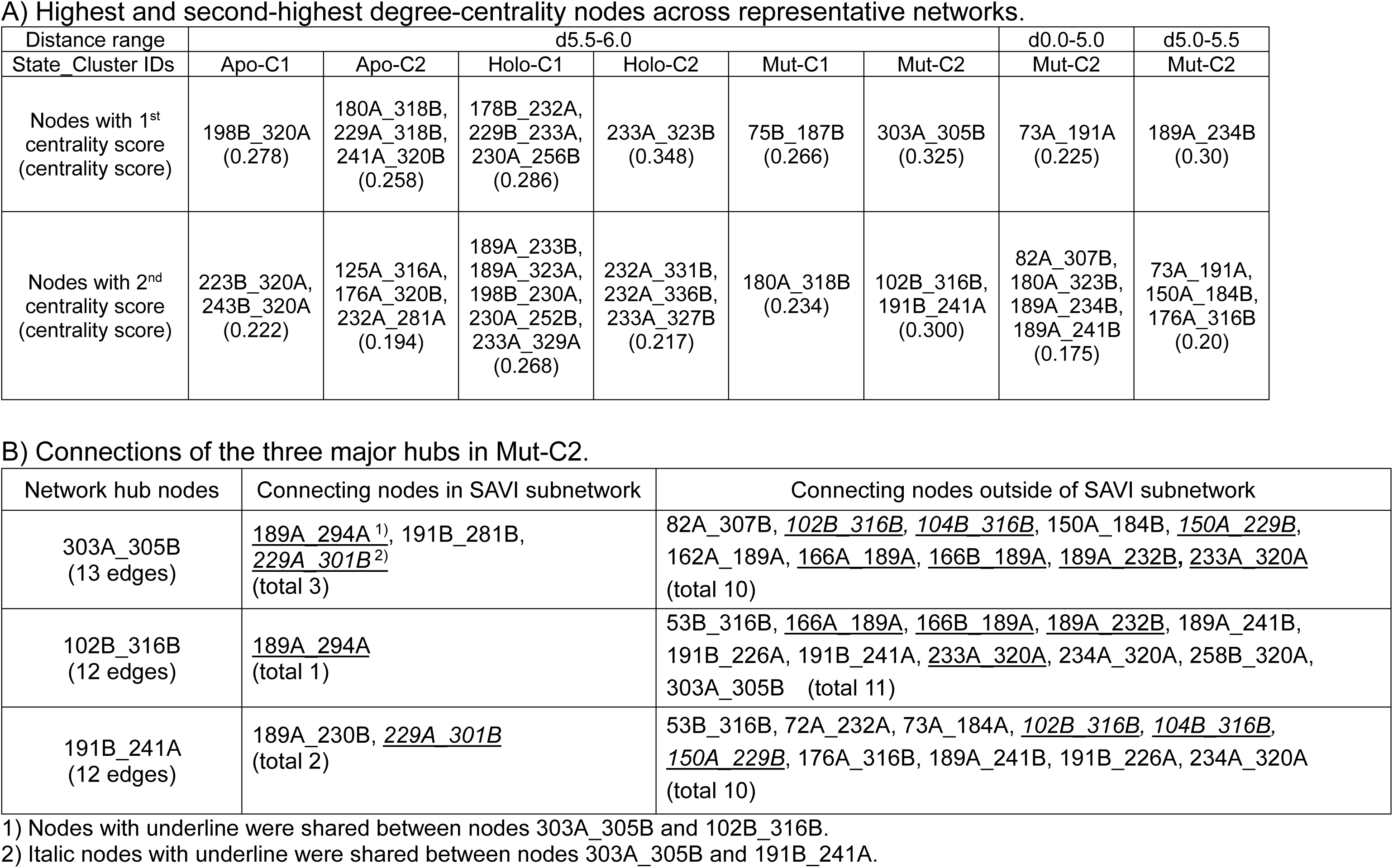
Top degree-centrality nodes and network hubs. A) Highest and second-highest degree-centrality nodes across representative networks. B) Connections of the three major hubs in Mut-C2. Degree centrality was calculated on the full graph before removal of isolated nodes or small disconnected components. For Mut-C2 in d5.5–6.0, the full input graph contained 41 nodes, whereas the reported major component contained 31; the degree centrality of 303A_305B was therefore 13/(41 - 1) = 0.325.

## Results

### Latent representations capture residue-pair fluctuation patterns

From analyzed MD trajectories, each residue pair was represented by an 1800-point time series of mean pairwise distances between side-chain heavy atoms extracted from coordinate-averaged structures in the 100– 1000 ns analysis window. Training the Conv1D-AE on apo trajectories produced a compact latent representation for each residue pair. The resulting 25-dimensional latent vectors summarize temporal patterns of residue-pair distance fluctuations and provide a common representation for comparing apo, holo, and mutant dynamics.

### Hierarchical clustering resolves distinct residue-pair fluctuation patterns

Agglomerative clustering using Mahalanobis distance placed the vast majority of residue pairs in a numerically dominant background cluster (>50,000 pairs), representing the common latent fluctuation behavior within each state. Nine low-population minor clusters (<100 pairs each) captured distinct fluctuation patterns outside this background population. We focused on the two largest minor clusters in each state because together they accounted for more than half of all non-background residue pairs (Table 1).

The sizes of these top minor clusters differed across states. In the apo state, the two clusters contained 66 and 38 pairs, respectively. In the holo state they contained 96 and 46 pairs, and in the mutant state they contained 82 and 58 pairs (Table 1). Thus, both perturbed states contained larger non-background populations than the apo state, while the composition of those populations differed substantially between ligand-bound and mutant conditions.

### State-specific cluster composition highlights distinct dominant regions

The top minor clusters showed markedly different regional compositions between states. In the holo state, both clusters were dominated by lid-containing pairs (≥85% of pairs; Table 1), indicating that ligand binding concentrates coordinated dynamics around the ligand-capping region and neighboring LBD elements. In the apo state, by contrast, the peripheral-LBD region (a3-b5/b5-a4/a4) contributed the majority of pairs in the two largest minor clusters (approximately 75%), with fewer lid- and a1b1-containing pairs (Table 1).

The SAVI-associated multi-mutant model displayed a distinct pattern. Although peripheral-LBD pairs remained substantial in Mut-C1, the a1b1 region was enriched relative to the apo and holo states. In particular, Mut-C2 contained a1b1 residues in approximately 40% of residue pairs, and these residues were paired not only with lid and peripheral-LBD regions but also with mutation-associated sites and other regions, including the transmembrane region (Table 1). This distribution suggests that the mutant state redistributes coordinated fluctuations toward an a1b1-centered, cross-region organization rather than reproducing the lid-dominated ligand-bound pattern.

### Network topology differs between ligand activation and SAVI-associated mutation

We translated the selected residue-pair sets into networks to examine how coordinated fluctuation signatures organize at a higher level. Nodes represent residue pairs, and edges connect nodes with similar latent fluctuation signatures within the three Mahalanobis-distance bands. We compared the two largest minor clusters in each state and paid particular attention to networks containing nodes involving residues 281 and/or 284 (Table 2). Figure 3 shows representative ligand-bound and mutant networks in the d5.5–6.0 band, with full-size networks and additional bands provided in Supplementary Figure 1.

**Figure 3.**
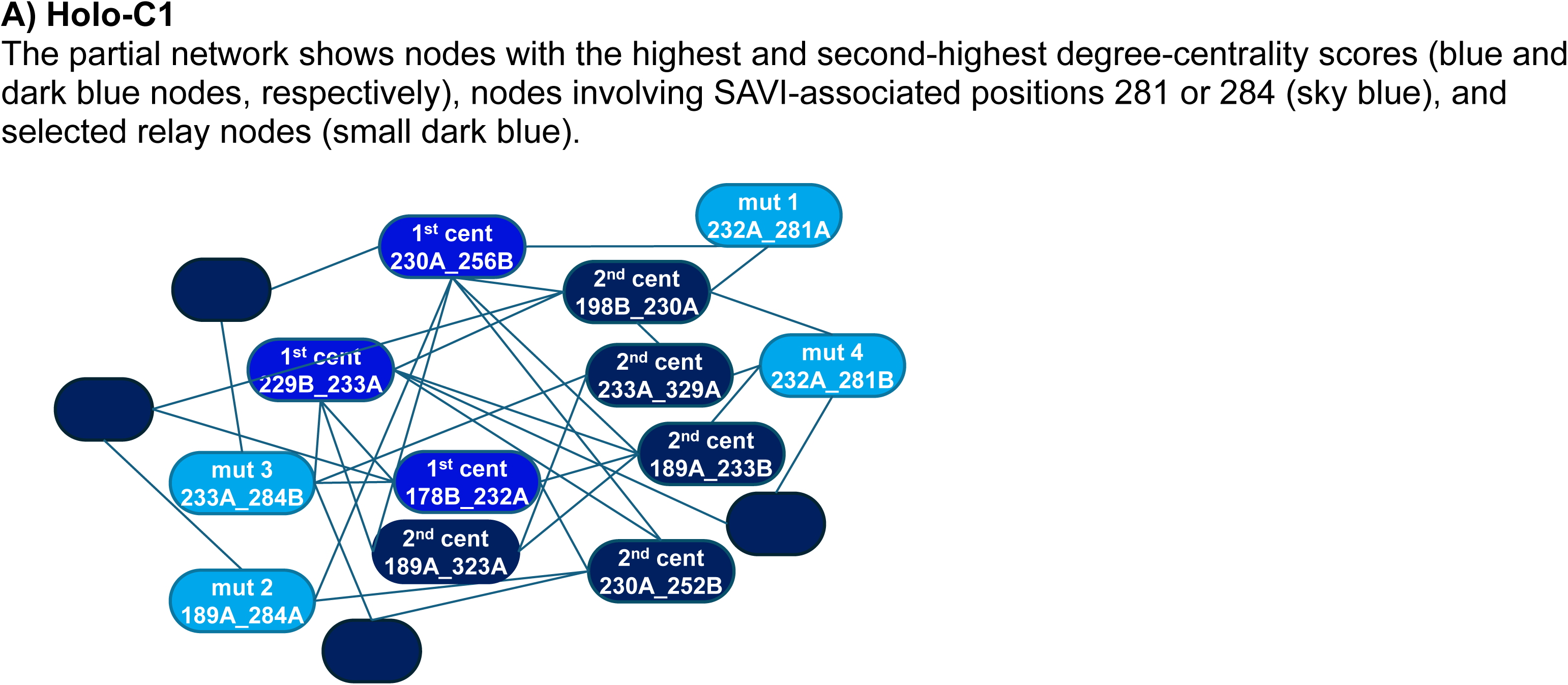
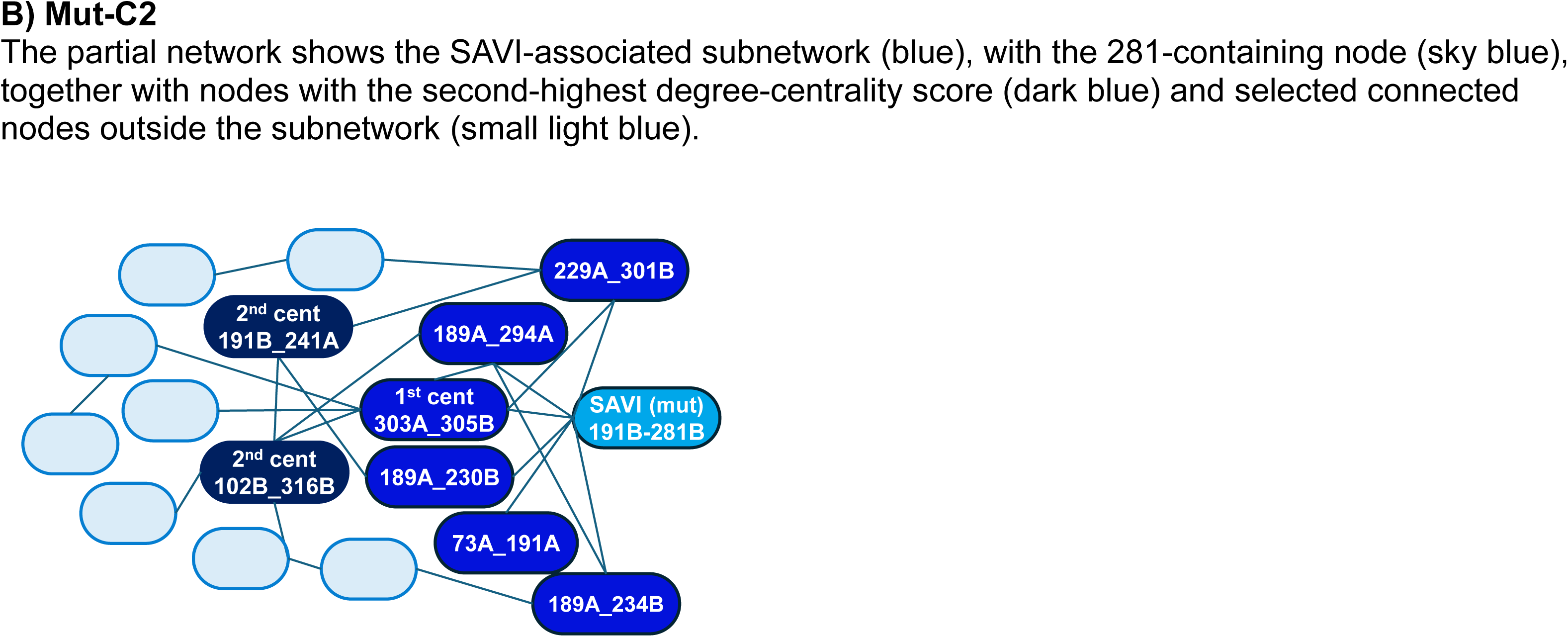
Representative latent-feature networks in the d5.5–6.0 Mahalanobis-distance band. Representative networks are shown for Holo-C1 (Figure 3A) and Mut-C2 (Figure 3B) to highlight the contrast between ligand-driven lid-centered organization and mutation-associated a1b1-enriched coupling. Nodes represent residue pairs and edges connect nodes whose latent vectors fall within the d5.5–6.0 Mahalanobis-distance band. Full-size network views for apo, holo, and mutant states, together with additional Mut-C2 similarity bands, are provided in Supplementary Figure 1.

### Apo networks are peripheral-LBD-dominated and sparsely involve SAVI hotspot residues

In the apo state, Apo-C1 formed relatively large networks dominated by peripheral-LBD nodes and lacked nodes involving residues 281/284 (Table 2A). Apo-C2 contained two residue-pair nodes involving wild-type residue 281: 232A_281A and 281B_336A. Both nodes were present in the reported major connected components for d0.0–5.0 and d5.5–6.0, whereas only 281B_336A was retained in the reported component for d5.0–5.5 (Table 2A).

In Apo-C2, the d5.5–6.0 network contained 25 nodes and 51 edges and was heavily enriched in peripheral-LBD nodes (92%), with lid and a1b1 nodes contributing 52% and 16%, respectively (Table 2A). The highest-centrality nodes involved peripheral-LBD residues (180A_318B, 229A_318B, and 241A_320B; Table 3A). Both wild-type 281-containing nodes were present, but neither belonged to the highest-degree-centrality group. Overall, the apo network was dominated by peripheral LBD-containing pairs, with residue 281 represented by a small number of nodes.

### Holo networks show lid-centered organization

Both selected clusters in the ligand-bound dataset produced lid-dominated networks. In Holo-C1, the major connected component in the d5.5–6.0 band contained 49 nodes and 192 edges (Table 2B). Nearly all nodes contained lid residues (93.9%, 46 nodes), consistent with strong coordination among ligand-responsive LBD elements. Holo-C2 also produced a smaller lid-rich network (20 nodes and 31 edges in d5.5–6.0; Table 2B), indicating that the two largest minor clusters capture distinct modes within the ligand-driven response.

Multiple nodes involving wild-type residues 281/284 were present in the holo networks, frequently as lid-hotspot pairs (Table 2B). However, centrality analysis showed that the highest-centrality nodes in Holo-C1 and Holo-C2 were primarily lid- and LBD-containing pairs (e.g., 178B_232A, 229B_233A, and 230A_256B in Holo-C1, and 233A_323B in Holo-C2). In contrast, 281/284-containing nodes had comparatively lower centrality and did not organize the network centers (Table 3; Figure 3A and Supplementary Figure 1B). Thus, in the analyzed ligand-bound dataset, the wild-type counterparts of SAVI hotspot residues participate in lid-rich networks without occupying the highest degree-centrality ranks, contrasting with the organization of the selected mutation-associated network.

### SAVI-associated mutant networks show a1b1-centered reorganization and cross-region coupling involving residue 281

The mutant model displayed two contrasting behaviors across its two largest minor clusters. Mut-C1 formed relatively large networks dominated by peripheral-LBD nodes and did not produce a major connected component containing residues 281/284 within the analyzed d0.0–6.0 range (Table 2C). In contrast, Mut-C2 exhibited a distinctive a1b1-enriched topology: in all three Mahalanobis-distance bands, more than half of the connected nodes contained a1b1 residues (Table 2C). Mut-C2 also contained nodes involving non-281/284 mutation-associated sites, including residues 72 and 166, indicating distributed coupling between mutation-associated positions and core structural regions.

In Mut-C2, the node 191B_281B was present in the reported major connected component for d5.5–6.0, where it and its six directly connected neighbors defined the SAVI-associated subnetwork (Figure 3B and Supplementary Figure 1C). The major connected component in this band contained 31 nodes and 101 edges; 17 (54.8%), 13 (41.9%), and 12 (38.7%) nodes contained a1b1, lid, and peripheral LBD residues, respectively (Table 2C). To compare the fluctuation relationships within this subnetwork, we examined the same six neighboring residue-pair nodes in d0.0–5.0 and d5.0–5.5. Several were directly connected to one another, whereas others were linked through relay nodes such as 82A_307B and 176A_316B (Supplementary Figure 1C). In contrast, 191B_281B was absent from the major connected components reported for these two bands (Table 2C). The band-wise comparison characterizes these residue pairs at different levels of latent-feature similarity.

To characterize connectivity within Mut-C2, we ranked nodes by degree centrality. The top hub was 303A_305B, with 13 connections and a degree centrality of 0.325 (Table 3A). This cross-protomer a3-b5 loop pair was directly connected to three nodes within the SAVI-associated subnetwork (189A_294A, 191B_281B, and 229A_301B) and 10 nodes outside it, including 82A_307B and 102B_316B (Table 3B). The next-highest centrality nodes, 102B_316B and 191B_241A, each had 12 connections (degree centrality 0.300; Table 3A) and lay outside the SAVI-associated subnetwork. Their overlapping neighborhoods with 303A_305B place the 281-containing subnetwork within a broader network of residue-pair fluctuation relationships.

Mapping these residue-pair nodes onto the STING structure provides a spatial interpretation of the network (Figure 4). The hub 303A_305B comprises residues in the a3-b5 loops of different protomers. Within the same subnetwork, 189A_294A pairs the a1b1 loop with the a3 helix, and 229A_301B pairs the lid region with the a3 helix. The subnetwork also contains a1b1–lid pairs (189A_230B and 189A_234B) and a TM2–TM3 linker–a1b1 pair (73A_191A). Beyond this subnetwork, 191B_241A pairs a1b1 and lid residues, whereas 102B_316B pairs the TM3 region with the peripheral LBD. Together, these residue-pair nodes map the network across the two protomers, ligand-responsive LBD elements, and membrane-proximal regions.

**Figure 4.**
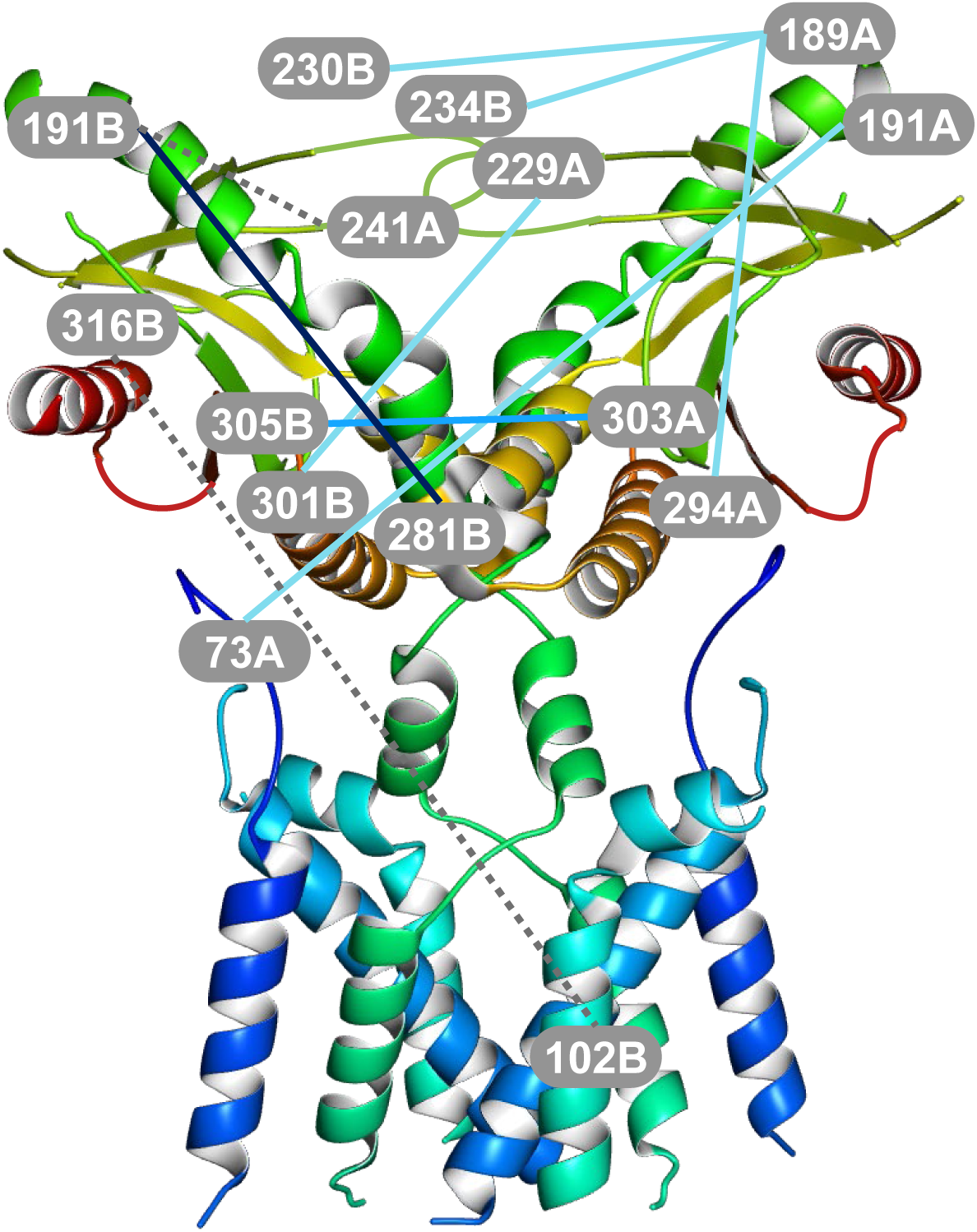
Structural mapping of network paths from the 281-containing node in Mut-C2 in d5.5–6.0. The dark-blue line denotes the 281-containing node 191B_281B. Blue and light-blue lines denote residue-pair nodes directly connected with 191B_281B, including the highest-centrality hub 303A_305B (blue line). Gray dotted lines indicate additional high-centrality nodes outside the SAVI-associated subnetwork, including 102B_316B and 191B_241A. The network is mapped onto the apo STING dimer structure to illustrate the spatial distribution of the participating regions.

Compared with the apo and holo networks (Supplementary Figure 2), the SAVI-associated multi-mutant network therefore reflects neither a peripheral-LBD-dominated apo-like regime nor a lid-dominated holo-like regime. Instead, it is characterized by a1b1 enrichment, cross-protomer hubs, and participation of residue 281 in a structurally distributed subnetwork, supporting a distinct mutation-associated dynamic organization.

## Discussion

This study presents a residue-pair-centric strategy that combines deep representation learning and network analysis to characterize protein dynamics, with STING as an application for comparing apo, ligand-bound, and SAVI-associated multi-mutant conditions. By compressing pairwise distance trajectories into latent vectors and clustering them to isolate non-background behaviors, we obtained compact residue-pair sets suitable for network analysis. The resulting networks provide a mesoscale representation of coordinated fluctuation signatures that is complementary to contact-based maps and static structural comparisons.

The recurrent emergence of a >50,000-pair background population in all three independently clustered states suggests that most residue-pair trajectories occupy a broad scaffold-level fluctuation repertoire that remains numerically dominant across STING conditions. Because clustering was performed separately for each state, this observation does not imply one-to-one identity of the background clusters at the residue-pair level. Rather, it indicates that state-dependent differences are concentrated in comparatively small, low-population sectors of latent-feature space, which are the focus of the present analysis.

The STING application illustrates how the workflow distinguishes ligand-associated and mutation-associated fluctuation patterns. In the ligand-bound dataset, the selected networks are dominated by lid-containing pairs and neighboring LBD elements. Wild-type 281/284-containing nodes also participate in these networks, whereas the highest-degree-centrality nodes primarily involve lid and neighboring LBD residues.

In contrast, Mut-C2 is a1b1-enriched and contains cross-region residue pairs involving the lid, peripheral LBD, and membrane-proximal elements. The 191B_281B node belongs to the reported d5.5–6.0 component, whereas several of its neighboring nodes are mutually connected in d0.0–5.0 or d5.0–5.5. This pattern associates residue 281 with a network of fluctuation relationships distributed across different parts of STING. Structural mapping places these relationships in the context of the dimer, suggesting a mode of dynamic coupling distinct from the lid-centered pattern in the ligand-bound model.

The structural contexts of the major hubs reinforce this interpretation. Enrichment of a1b1-containing nodes places a candidate coordination center adjacent to the ligand-responsive lid, while the cross-protomer hub 303A_305B links this organization to the LBD periphery and dimer interface. The TM2-TM3-linker/a1b1 pair 73A_191A together with the additional hubs 191B_241A and 102B_316B extend the same network toward lid and membrane-proximal regions, respectively. These observations suggest a distributed, dimer-spanning organization in the selected mutant network, contrasting with the more localized lid-centered organization of the ligand-bound dataset.

More broadly, the latent-space residue-pair network framework provides a practical way to summarize long MD trajectories by identifying which regions become dynamically coordinated and which residue-pair nodes act as hubs or bridges in each state. By focusing on residue-pair dynamics rather than residue-wise metrics, the approach can capture non-local and cross-region coupling signatures that may be missed by conventional summary measures, providing a general route from large MD-derived time-series datasets to interpretable dynamic-network hypotheses.

The present application uses one microsecond-scale trajectory per condition to demonstrate the complete analysis sequence, from distance time series to structural interpretation. The comparison characterizes the three analyzed datasets; application to independent trajectories would allow the consistency of these patterns across runs to be assessed.

Overall, the workflow distinguishes lid-centered fluctuation patterns in ligand-bound STING from a1b1-enriched, dimer-spanning patterns in the multi-mutant model. By linking MD-derived time series to interpretable residue-pair networks, it provides a practical framework for comparing protein dynamics across molecular conditions and developing structurally grounded hypotheses of dynamic allostery.

## Supporting Information

Full-size latent-feature networks for apo, holo, and SAVI-associated mutant states, including additional Mut-C2 Mahalanobis-distance bands; structural mapping of 281/284-containing network paths in apo and holo states; representative 159-residue list; clustering results (PDF).

## Data and Software Availability

The postprocessed protein-coordinate trajectories and corresponding GROMACS TPR files used for the reported analyses, standardized residue-pair distance datasets, trained autoencoder model, latent descriptors, clustering outputs, network-analysis data, and analysis scripts are available in Zenodo at https://doi.org/10.5281/zenodo.23232081 [24].

