## Supplementary material for "Latent-Space Residue-Pair Networks from Molecular Dynamics Distinguish Ligand-Activated and SAVI-Associated STING": Supporting_Information.pdf

#### Supplementary Figure 1:

#### Full-size latent-feature networks for apo, holo, and SAVI-associated mutant states, including additional Mut-C2 Mahalanobis-distance bands.

Nodes represent residue pairs, and edges connect nodes whose latent descriptors fall within the stated Mahalanobis-distance band. Smaller distances indicate greater similarity in the learned descriptors, not shorter physical inter-residue distances. The three bands select different edges from the same analyzed time series; they are neither successive time intervals nor cumulative cutoff networks. "Mutant node" denotes a node containing a residue position mutated in the multi-mutant model; the corresponding residues retain their wild-type identities in apo and holo.

A) Apo-C2 network in the d5.5-6.0 Mahalanobis-distance band.
The major connected component contains 25 nodes and 51 edges. Of these, 23 (92%) contain at least one peripheral LBD residue. Two nodes contain wild-type residue 281 (232A_281A and 281B_336A), but neither belongs to the highest-degree-centrality group. Overall, Apo-C2 is dominated by peripheral LBD-containing pairs, with limited representation of residue 281-containing nodes.

B-1) Holo-C1 network in the d5.5–6.0 Mahalanobis-distance band.
The major connected component contains 49 nodes and 192 edges; 46 nodes (93.9%) include at least one lid residue. Several nodes also contain wild-type residues at SAVI-associated positions. The highest degree-centrality ranks are occupied by the residue-pair nodes listed in Table 3A rather than by the 281/284-containing nodes. These results describe a predominantly lid-centered organization within the analyzed ligand-bound dataset.

B-2) Holo-C2 network in the d5.5-6.0 Mahalanobis-distance band.
The major connected component contains 20 nodes and 31 edges. It includes three nodes involving SAVI-associated hotspot positions paired with lid residues (147A_233A, 233A_281B, and 233A_284A). Nineteen of 20 nodes (95.0%) contain at least one lid residue, 14 (70.0%) contain a peripheral-LBD residue, and 3 (15.0%) contain an a1b1 residue, further supporting the strong lid-centered organization of the holo state.

C-1) Mut-C2 network in the d5.5–6.0 Mahalanobis-distance band.
The major connected component contains 31 nodes and 101 edges, including 191B_281B. Degree centrality was calculated on the full 41-node input graph: 303A_305B has 13 incident edges and centrality 0.325; 102B_316B and 191B_241A each have 12 incident edges and centrality 0.300. Within the 281-associated subnetwork, 303A_305B is directly connected to 191B_281B, 189A_294A, and 229A_301B (Table 3B). Structural mapping places 189A_294A between the a1b1 loop and a3 helix, and 229A_301B between the lid and a3 helix.

C-2) Mut-C2 network in the d0.0-5.0 Mahalanobis-distance band.
The major connected component contains 23 nodes and 47 edges and excludes nodes containing residues 281 or 284. Several residue-pair nodes that belong to the 281-associated subnetwork in the d5.5-6.0 graph are connected to one another in this higher-similarity graph, either directly or through relay nodes. This is a comparison between edge-distance bands derived from the same time series; it does not identify an earlier stage of subnetwork formation.

C-3) Mut-C2 network in the d5.0-5.5 Mahalanobis-distance band.
The major connected component contains 29 nodes and 63 edges and excludes nodes containing residues 281 or 284. Non-281 members of the subnetwork displayed in d5.5-6.0 show direct or relay-mediated relationships in this band. Together with the d0.0–5.0 graph, this comparison shows that these non-281 relationships occur in higher-similarity bands, whereas the displayed edges involving 191B_281B occur in d5.5–6.0.

### A) Apo-C2 network in the d5.5-6.0 Mahalanobis-distance band.

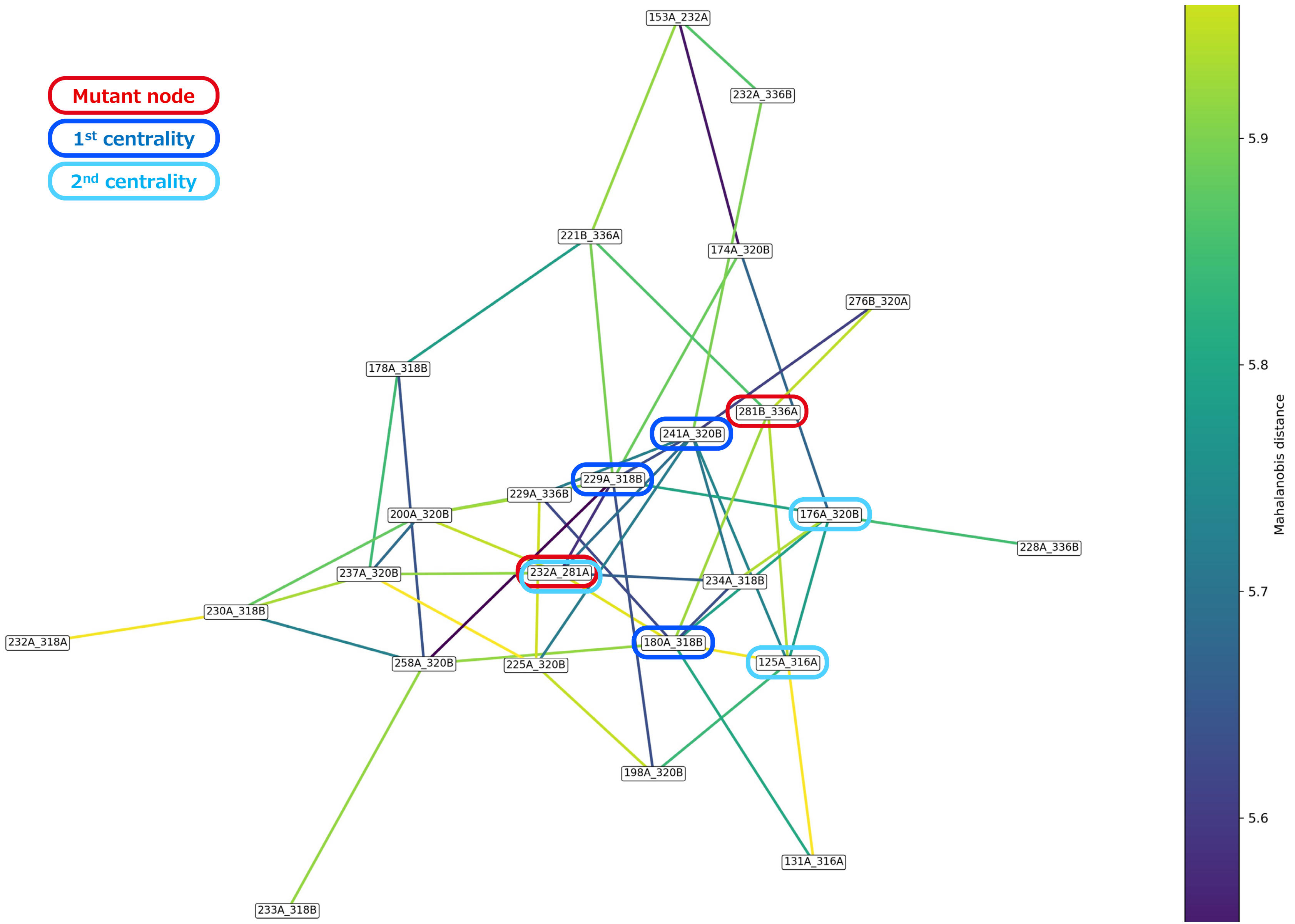

#### B-1) Holo-C1 network in the d5.5-6.0 Mahalanobis-distance band.

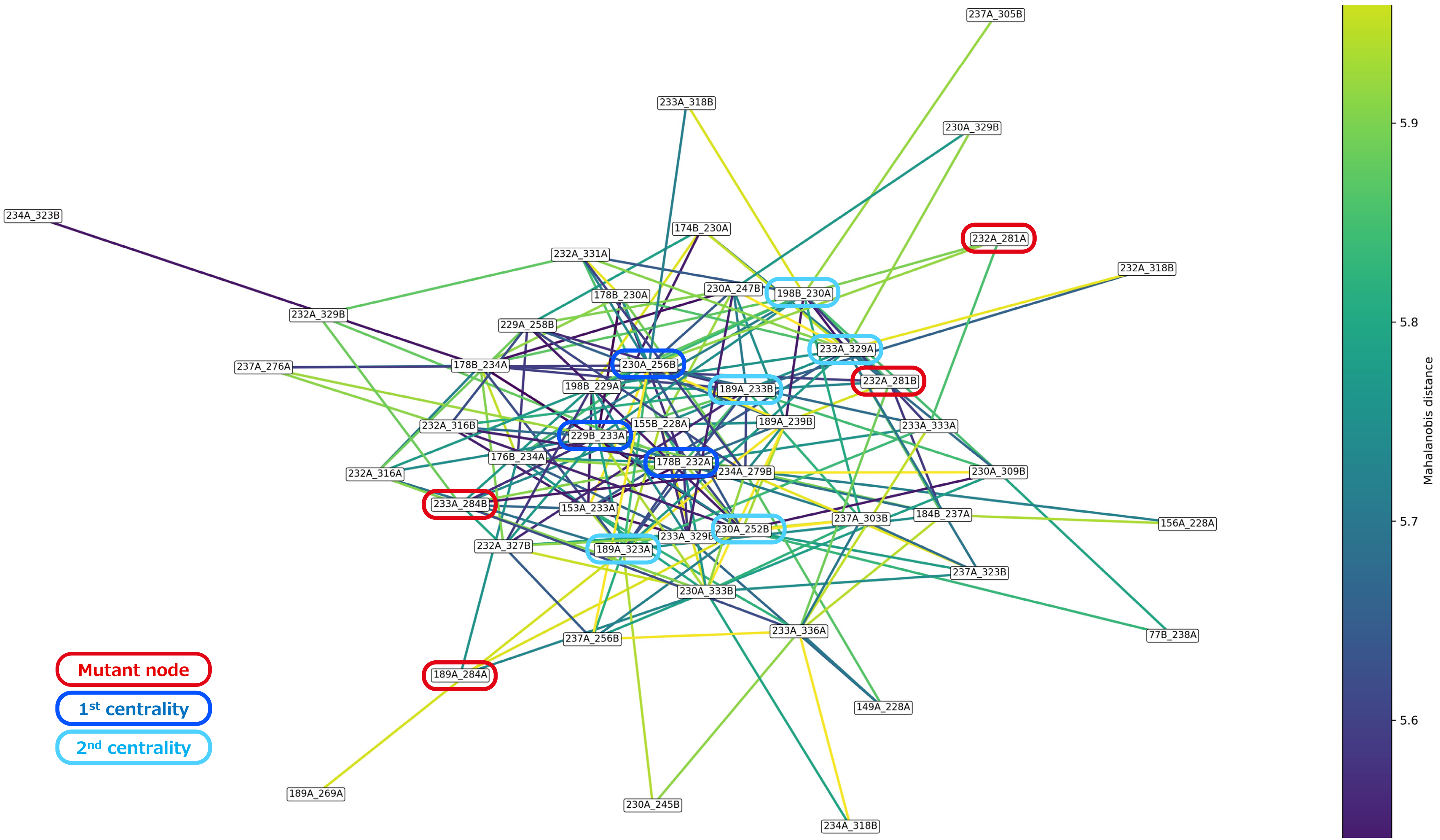

#### B-2) Holo-C2 network in the d5.5-6.0 Mahalanobis-distance band.

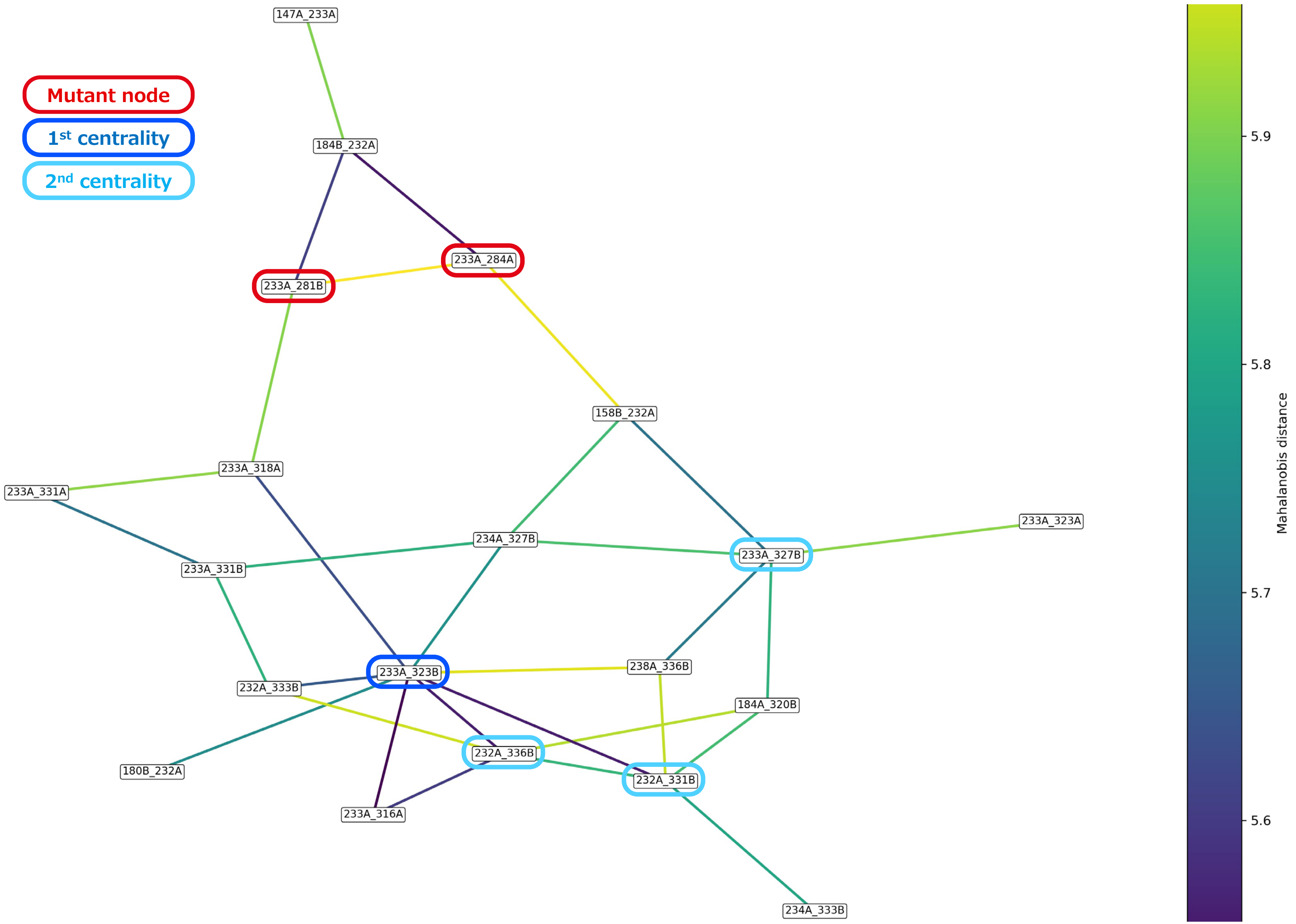

#### C-1) Mut-C2 network in the d5.5-6.0 Mahalanobis-distance band.

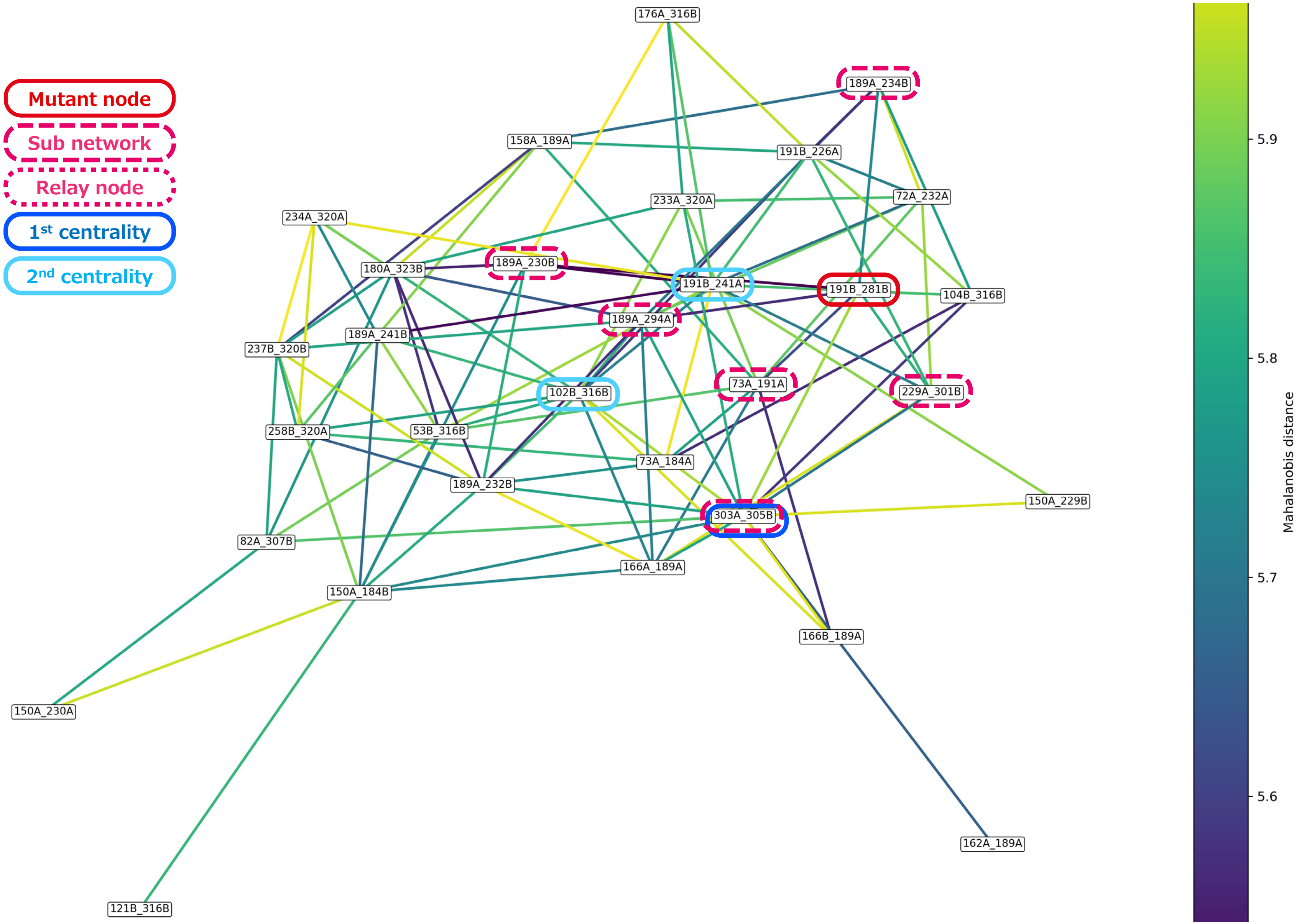

### C-2) Mut-C2 network in the d0.0-5.0 Mahalanobis-distance band.

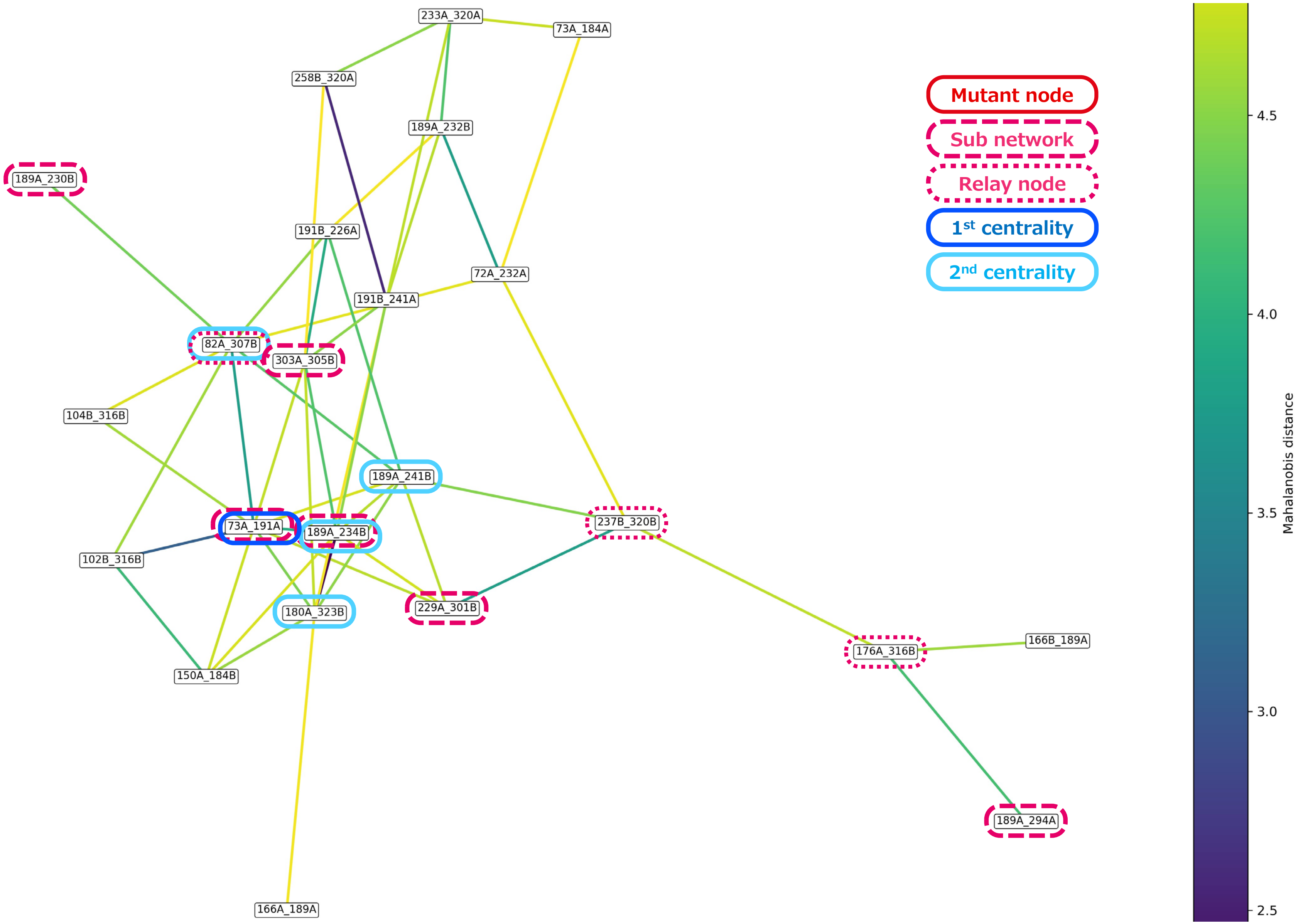

#### C-3) Mut-C2 network in the d5.0-5.5 Mahalanobis-distance band.

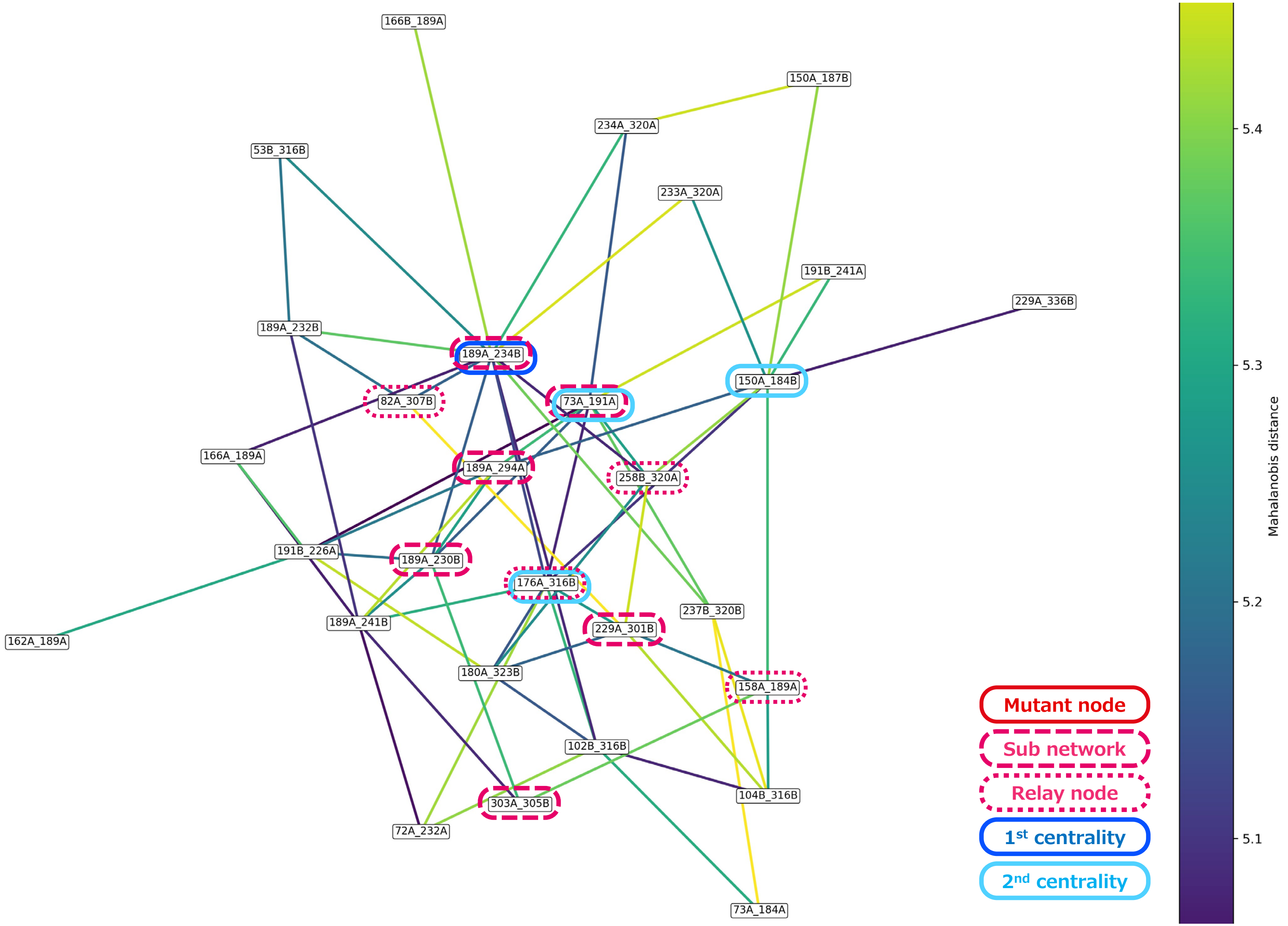

#### Supplementary Figure 2:

#### Structural mapping of 281/284-containing network paths in apo and holo states.

The dark-blue line identifies the residue-pair node containing residue 281 or 284. Light-blue lines denote residue-pair nodes directly connected to the indicated hotspot-containing node in the latent-feature network. These lines therefore summarize similarity relationships between time-dependent fluctuation signatures and should not be interpreted as direct physical contacts. Apo-C2 and the two holo networks (Holo-C1 and Holo-C2) are mapped onto the corresponding STING dimer structures.

#### A) Apo-C2 network in the d5.5–6.0 Mahalanobis-distance band.

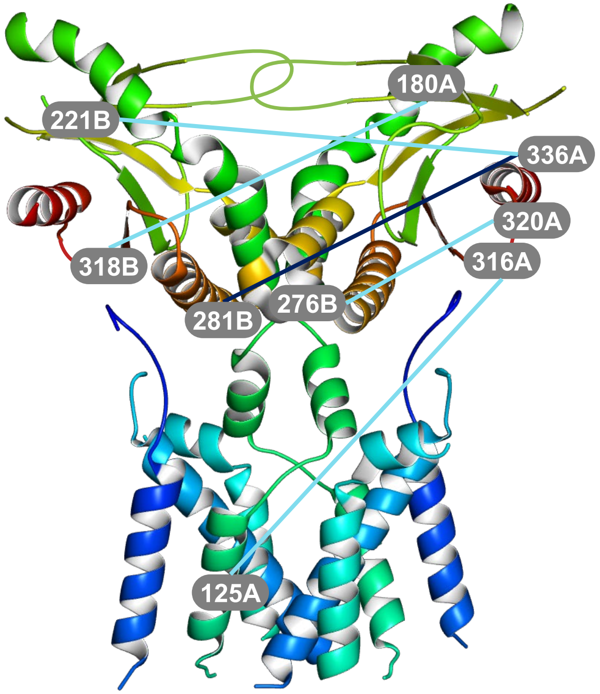


Paths from 281B_336A

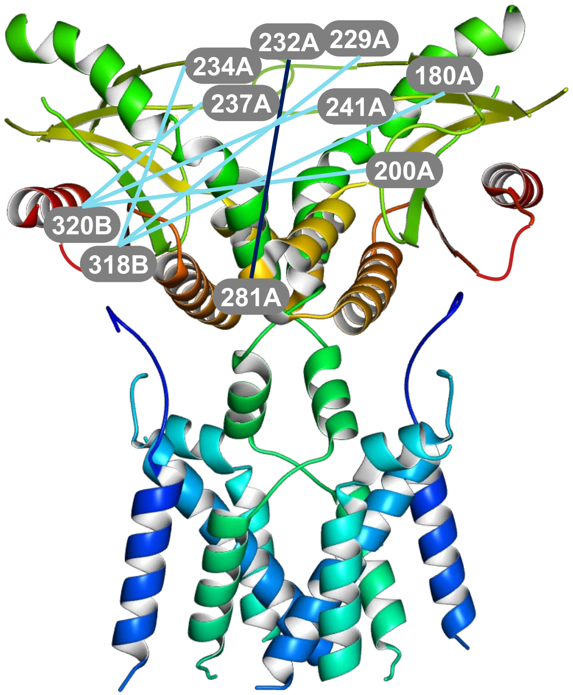


Paths from 232A_281A

#### B-1) Holo-C1 network in the d5.5–6.0 Mahalanobis-distance band.

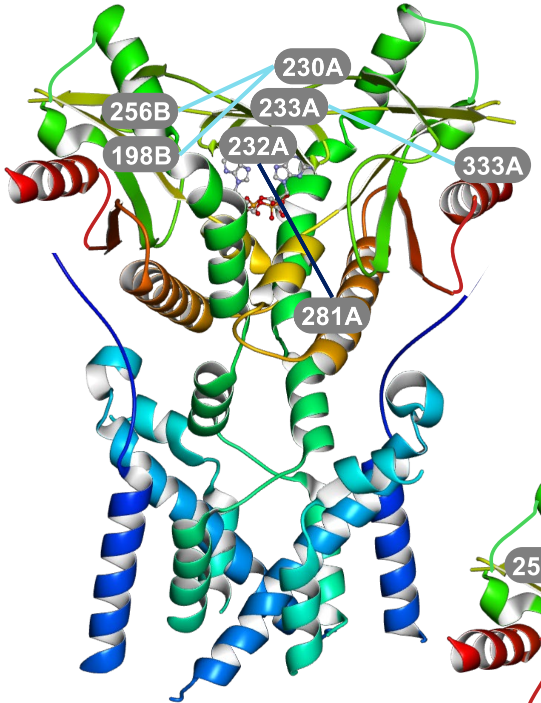


Paths from 232A_281A

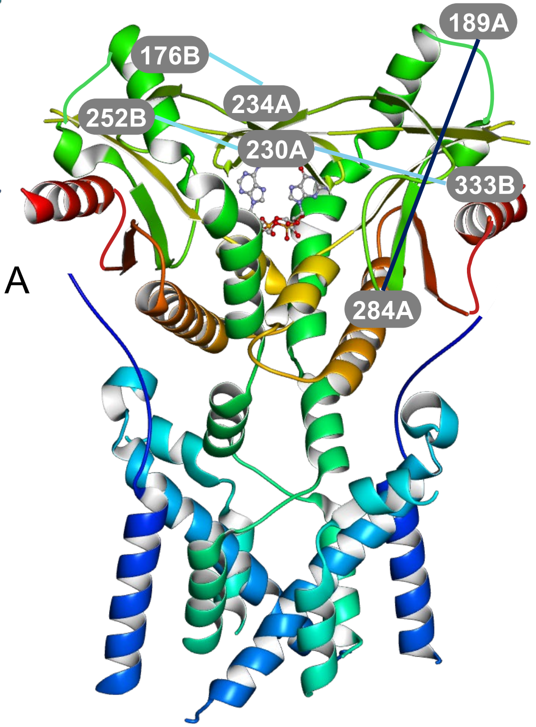


Paths from 189A_284A

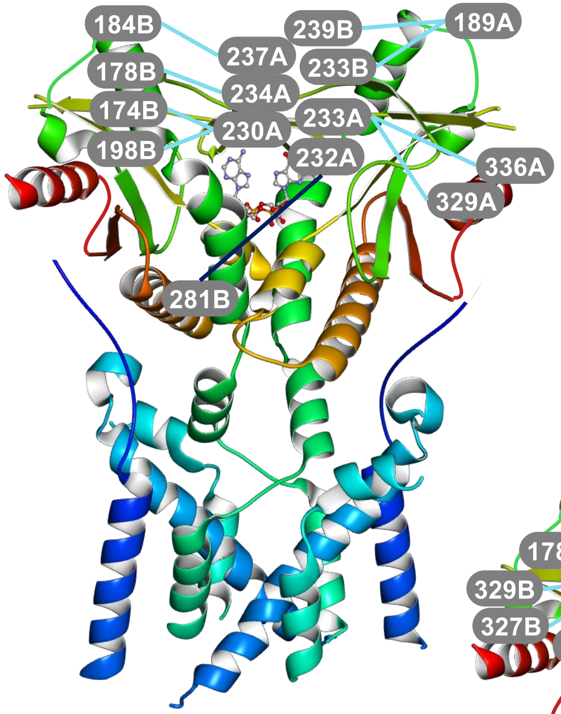


Paths from 232A_281B

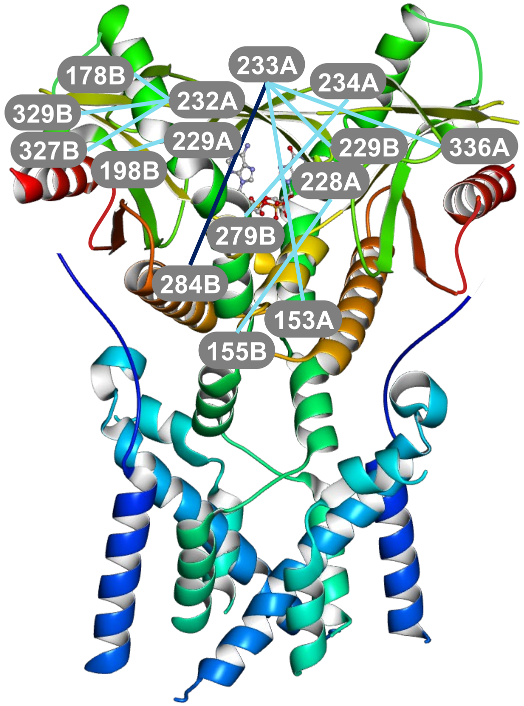


Paths from 233A_284B

**B-2) Holo-C2 network in the d5.5–6.0 Mahalanobis-distance band.**

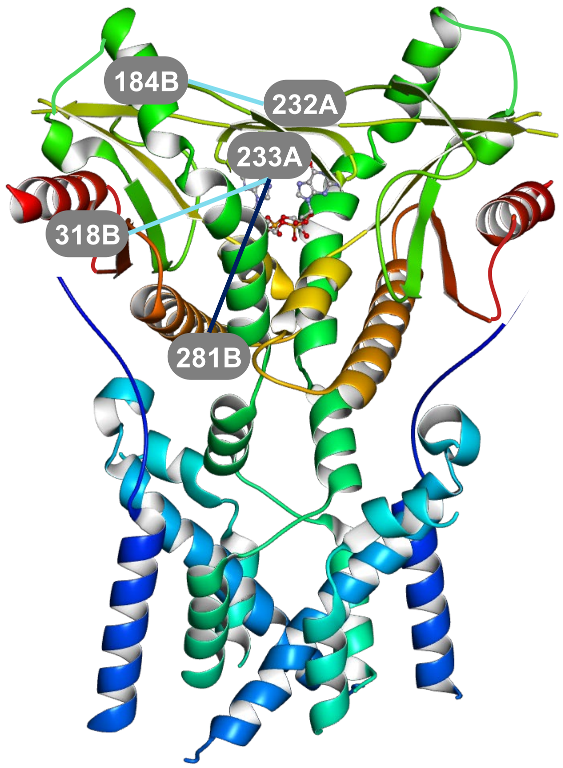


Paths from 233A_281B

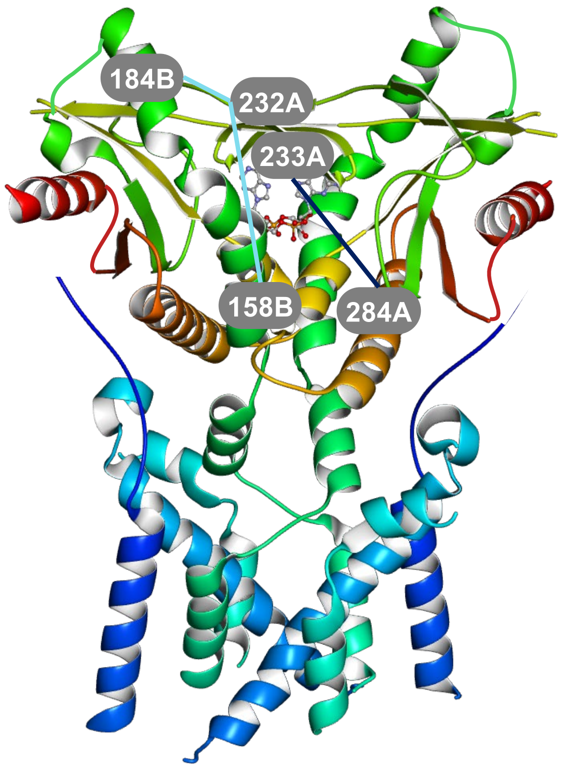


Paths from 233A_284A

#### Supplementary Table 1:
#### Representative 159 residues focused on in this study.

| Regions | Representative residues | #residues |
|---|---|---|
| TM1 helix (tm1: 18-34) | 18/20/22/24/26/28/30/32/34 | 9 |
| Loop between TM1 and TM2 (tm12: 35-44) | 36/38/40/43 | 4 |
| TM2 helix (tm2: 45-69) | 45/47/49/51/53/55/57/59/61/63/65/67/69 | 13 |
| TM2-3 linker region (tm23: 70-91) | 71/72(savi)/73/75/77/79/82/84/86/88/90 | 11 |
| TM3 helix (tm3: 92-106) | 92/94/96/98/100/102/104/106 | 8 |
| Loop between TM3 and TM4 (tm34: 107-116) | 108/110/112/115 | 4 |
| TM4 helix (tm4: 117-134) | 117/119/121/123/125/127/129/131/134 | 9 |
| Loop between TM4 and Connector helix (tm4cn: 135-140) | 136/139 | 2 |
| Connector helix (cnhx: 141-149) | 141/143/145/147(savi)/149 | 5 |
| Connector loop (cnlp: 150-156) | 150(oth)/152/153(wksv)/154(savi)/155(savi)/156 | 6 |
| A1_former helix (a1fm: 157-175) | 158(wksv)/160/162/164/166(savi)/168/170/172/174 | 9 |
| A1_latter helix (a1lt: 176 -187) | 176/178/180(ab1)/182(ab1)/184(ab1)/187(ab1) | 6 |
| Loop between A1 and B1 (a1b1: 188-197) | 189(ab1)/191(ab1)/193(ab1)/196(ab1) | 4 |
| B1 sheet (b1: 198-203) | 198/200/202 | 3 |
| Loop between B1 and B2 (b1b2: 204-218) | 204/206(savi)/207(wksv)/208/210/212/214/217 | 8 |
| B2 sheet (b2: 219-230) | 219/221/223/225/226/228(lid)/229(lid)/230(lid) | 8 |
| Loop between B2 and B3 (b2b3: 231-237) | 232/233/234/237(lid) | 4 |
| B3 sheet (b3: 238-249) | 238(lid)/239(lid)/241/243/245/247/249 | 7 |
| Loop between B3 and B4 (b3b4: 250-251) | - | 0 |
| B4 sheet (b4: 252-261) | 252/254/256/258/261 | 5 |
| Loop between B4 and a2 (b4a2: 262) | - | 0 |
| A2 helix (a2: 263-272) | 263/265/267/269/272 | 5 |
| Loop between A2 and A3 (a2a3: 273-280) | 274/276/279(wksv) | 3 |
| A3 helix (a3: 281-301) | 281(savi)/284(savi)/286/288/290/292/294/296/298/301 | 10 |
| Loop between A3 and B5 (a3b5: 302-308) | 303/305/307 | 3 |
| B5 sheet (b5: 309-314) | 309/311/314 | 3 |
| Loop between B5 and A4 (b5a4: 315-324) | 316/318/320/323 | 4 |
| A4 helix (a4: 325-336) | 325/327/329/331/333/336 | 6 |
| Total #residues focused on in this study | | 159 |

**Supplementary Table 2:**
**Clustering results.**

Numbers of residue pairs assigned to the ten clusters in each state. Each state contained one numerically dominant background cluster (>50,000 pairs) and nine lower-population clusters (<100 pairs each). The two largest minor clusters were selected for subsequent network analysis.

| | C0 (>50000, background cluster) | C1 (largest minor cluster) | C2 (2nd largest minor cluster) | C3 | C4 | C5 | C6 | C7 | C8 | C9 | C1-C9 total |
|---|---|---|---|---|---|---|---|---|---|---|---|
| Apo | 50,273 | 66 | 38 | 13 | 4 | 4 | 2 | 1 | 1 | 1 | 130 |
| Holo | 50,158 | 96 | 46 | 29 | 26 | 20 | 15 | 6 | 4 | 3 | 245 |
| SAVI-associated mutant | 50,211 | 82 | 58 | 16 | 11 | 11 | 5 | 5 | 3 | 1 | 192 |
